# Extracellular Forces Cause the Nucleus to Deform in a Highly Controlled Anisotropic Manner

**DOI:** 10.1101/027888

**Authors:** Kristina Haase, Joan K. L. Macadangdang, Claire H. Edrington, Charles M. Cuerrier, Sebastian Hadjiantoniou, James L. Harden, Ilona S. Skerjanc, Andrew E. Pelling

**Author notes:** Correspondence To: Andrew E. Pelling, 150 Louis Pasteur, MacDonald Hall, University of Ottawa, Ottawa, ON K1N 6N5, Canada, Web: http://www.pellinglab.net.

## Abstract

Physical forces arising in the extra-cellular environment have a profound impact on cell fate and gene regulation; however the underlying biophysical mechanisms that control this sensitivity remain elusive. It is hypothesized that gene expression may be influenced by the physical deformation of the nucleus in response to force. Here, using 3T3s as a model, we demonstrate that extra-cellular forces cause cell nuclei to rapidly deform (< 1 s) preferentially along their shorter nuclear axis, in an anisotropic manner. Nuclear anisotropy is shown to be regulated by the cytoskeleton within intact cells, with actin and microtubules resistant to orthonormal strains. Importantly, nuclear anisotropy is intrinsic, and observed in isolated nuclei. The sensitivity of this behaviour is influenced by chromatin organization and lamin-A expression. An anisotropic response to force was also highly conserved amongst an array of examined nuclei from differentiated and undifferentiated cell types. Although the functional purpose of this conserved material property remains elusive, it may provide a mechanism through which mechanical cues in the microenvironment are rapidly transmitted to the genome.

## Introduction

Mechanical forces transmitted through the cell directly affect nuclear shape and function, have been implicated in altered gene expression^1,2^, and affect numerous processes at the cellular level^3,4^. Such forces result in internal remodelling of the nuclear cytoarchitecture and chromatin^3,5,6^, leading to alterations in transcriptional activity^7,8^. How nuclei respond to physical cues depends on their inherent material properties, which alone direct diverse biological functions. Mechanosensitive proteins, such as lamins, are known to control and regulate these properties by physically coupling the inner nucleus with the cell’s cytoskeleton, focal adhesions and integrins^3,5^. Mutations in these proteins result in a number of diseased states, and manifest in misshapen nuclei and increased nuclear fragility, particularly under applied strain^9-11^. The importance of nuclear mechanics is evidenced during differentiation and development wherein mechanical forces are necessary^12,13^ to direct deformable undifferentiated stem cells into committed cell types^14,15^. Matrix stiffness also influences lamin-A expression^16^ thereby governing nuclear resistance to force^17-19^. Clearly, characterizing how the nucleus deforms and remodels in response to force is of critical importance.

A large body of research involves examination of isolated nuclei^10,20,21^, as well as characterization of nuclei within their cyto-architecture^9,14,15,22,23^. Importantly, nuclei, and their constituents are often denoted as homogeneous, or isotropic materials^9,14,20,24,25^, despite displaying anisotropic material properties during physical perturbation^15,22,23^. Laser ablation studies have shown that the disruption of heterochromatin nodes causes elliptical nuclei to undergo anisotropic shrinkage in which they collapse significantly more along their minor axis as opposed to their major axis^15^. Planar stretching of cell monolayers also reveals an inherent mechanical anisotropy^23^. Moreover, whole-cell compression with a custom-built indenter revealed strain anisotropy in mouse embryonic fibroblast (MEF) nuclei, an observation that diminished in lamin A/C deficient cells^22^. These studies demonstrated that nuclear prestress governs nuclear shape, while cytoskeletal organization governs nuclear deformation in response to mechanical stress^5,11,15^. Whether or not the observed mechanical anisotropy is a widespread phenomenon amongst cells remains an open question. Moreover, if an intrinsic nuclear anisotropy exists, how is it regulated? Do both cytoskeletal and nuclear components influence this behavior simultaneously? It seems likely that the cytoskeleton plays a role in directing anisotropy of nuclear deformations since it acts as a direct link to external force. However, nuclear components such as lamins or chromatin organization may also inherently influence nuclear behaviour.

Our main objective in this study is to characterize the anisotropic deformation of nuclei within living cells in response to controlled extra-cellular forces. By measuring nuclear strains and defining a quantitative measure of anisotropy, we demonstrate that nuclear anisotropy is prominent in NIH 3T3 fibroblasts, and highly conserved in a variety of cell types. Using 3T3s as a model, we investigate the role of cytoskeletal components in nuclear deformation. We also discuss the implications of chromatin organization and lamin-A expression in directing nuclear mechanics. Nuclear anisotropy may provide a direct mechanism through which extra-cellular forces are controllably transmitted to the genome, thus the nucleus itself may act as a mechanosensor in the cell, as recently proposed^4^.

## Results

### Nuclear deformations reveal a mechanical anisotropy

Nuclear deformations were examined by applying a constant force above the nuclei of 3T3 fibroblasts using a pyramidal atomic force microscope (AFM) tip, while simultaneously acquiring time-lapse images (Fig. 1a,b) using laser scanning confocal microscopy (LSCM), in a manner previously described^6,26^. Employing the nuclear dye Hoechst 33342 enabled us to visualize nuclear area during the deformation as the tip was rapidly approached (10 μm/s) (Fig. 1a). The confocal acquisition plane was set in the middle of the nucleus (region of greatest fluorescence intensity), and images were collected at a rate of 4 frames/s. In order to avoid any artifacts due to focal drift, we analyzed the first 5 s of data. Longer durations were also examined (∼20 s); however no significant difference was observed in the total deformation. Confocal volume images demonstrated disk-shaped nuclei with an average height of ∼10 μm (n=5) and a conserved nuclear volume during the 10 nN deformation (mean volume insignificantly decreased by ∼5%). Image analysis allowed us to measure the change in nuclear area (Fig. 1c), and quantify strain (ε) along the major and minor axes of the nucleus as a function of time (Fig. 1d,e) as we described previously^23^.

**Figure 1.**
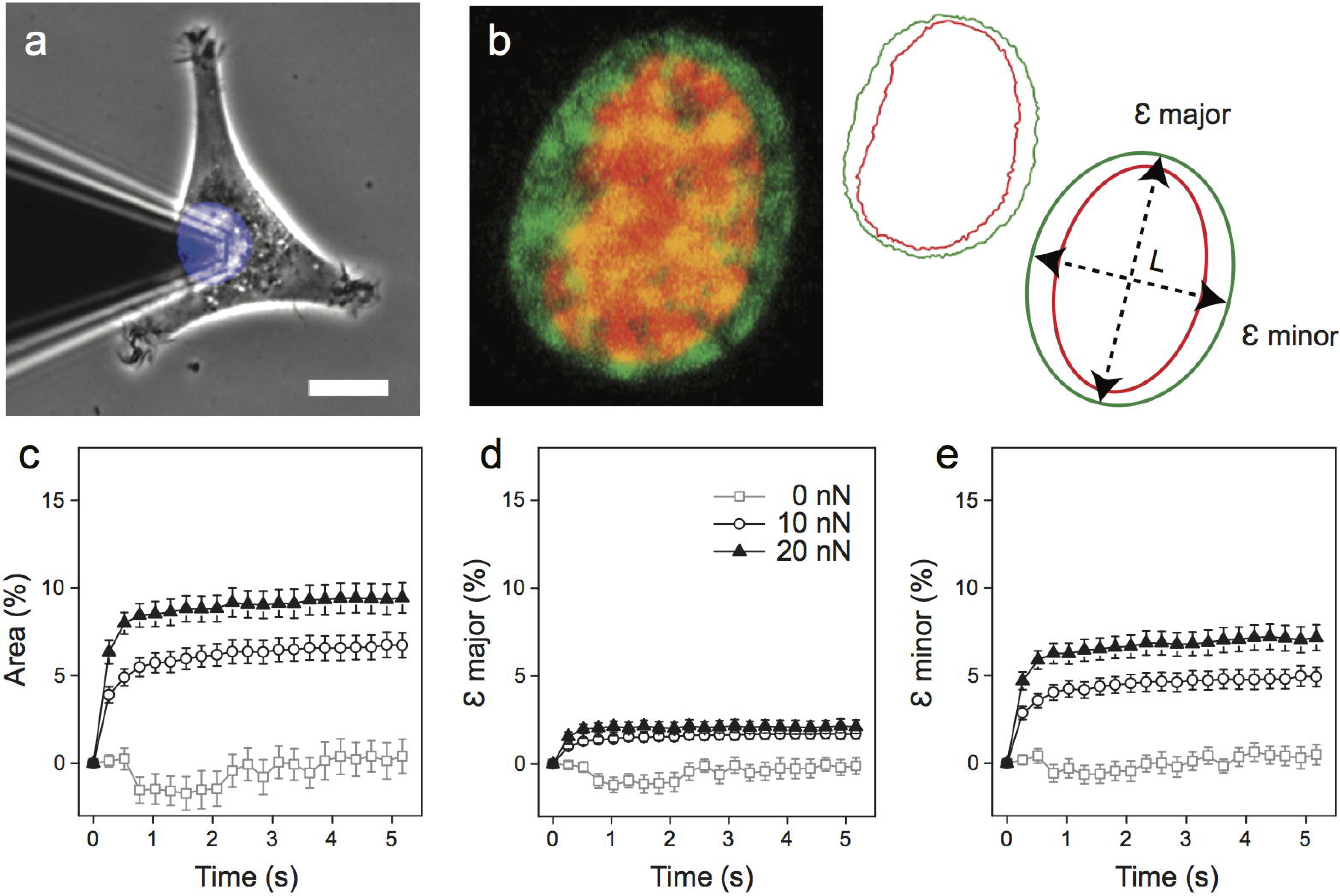
Extracellular forces applied to cell nuclei reveal anisotropic deformations. (a) Force application over the central nuclear region was carried out using AFM while simultaneously tracking nuclear deformation using LSCM. Scale bar is 10 μm. (b). Process of strain measurement. Nuclei stained with Hoechst 33342 allowed for observation of nuclear deformation. Shown is a 3T3 fibroblast nucleus prior to loading (red), and following force application (green), image is 12 μm wide. Nuclear outlines were extracted: before (red) and during (green) the deformation. Outlines were fit to an ellipse in order to extract the change in area and strain along the major (εmajor) and minor (εminor) axes over time, where ε = ΔL/L for each respective axis. (c) Area expansion following 5 s of 0 nN (□) n=15, 10 nN (○) n=32, and 20 nN (▴) n=29. (d) Percent strain over time in the major axis, and (e) the minor nuclear axis. Axial strains reveal a clear mechanical anisotropy. Shown is mean ± s.e.m.

Within 1 s following force application, nuclear area rapidly expanded (Fig. 1c). A clear dependence of nuclear expansion on force magnitude is evident (between 10 nN and 20 nN, P < 0.02), with no change in the absence of an applied force (0 nN area at 0 and 5 s, P > 0.9). Nuclear area increased by 6.7 ± 0.7 % and 9.4 ± 0.9 % following 5 s of a 10 nN and 20 nN constant force, respectively. Importantly, nuclear deformation was anisotropic, with absolute deformation along the major axis significantly less than the minor axis (P < 1×^-6^, paired t-tests). This anisotropy is demonstrated by plots of the force dependent strain, 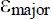 and 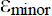, as a function of time (Fig. 1d,e). Following loading, 3T3 nuclei experienced strains of 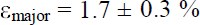 and 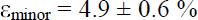 in response to 10 nN, and larger strains of 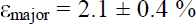 and 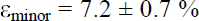 when exposed to 20 nN of force. The nucleus was confirmed not to move in the confocal volume due to mechanical loading, as demonstrated by stationary nucleoli during nuclear expansion (see Supplementary Fig. S1 a-c and Vid. S1). Force fluctuations were minimal during the deformation (s.d. = 0.18 nN) in relation to the setpoint force (10nN) over time (see Supplementary Fig. S1f).

To quantify the observed anisotropic strain dynamics, we fit the time dependent strain data to an expression that predicts deformation of a viscoelastic Kelvin-Voigt material^27^ (see Supplementary Methods). From the fits, we extracted plateau strains 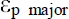 and 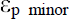, and characteristic deformation time constants 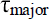 and 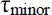. We then defined a dimensionless anisotropic stretch ratio 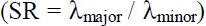, where the relative stretch (λ) can be approximated by 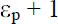, for each respective axis. A value of SR = 1 indicates isotropic deformation 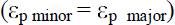, whereas SR < 1 implies anisotropic deformation, with 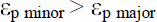. Accordingly, SR > 1 also implies anisotropy, but was rarely observed. Stretch (λ, an extension ratio) was used instead of engineering strain 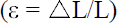 to avoid indeterminate ratios of small strains. Values of SR are generally > 0.85, since a large difference in strains between the nuclear axes would be required to grossly deviate from 1. For example, with 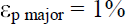 and 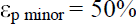, SR ≈ 0.67. Herein, mean strains were well under 20% for both axes. For instance, untreated 3T3s exposed to 10 nN of force resulted in plateau strains of approximately 2% and 5% in the major and minor axes, resulting in SR ≈ 0.971, which is significantly anisotropic (P < 1×10^−6^, one sample t-test compared to test mean for isotropy = 1).

Stretch ratios were independent of load magnitude (P > 0.05, two sample t-test) with mean SR = 0.971 ± 0.005 for 10 nN and SR = 0.956 ± 0.007 for 20 nN. Taken together, these results demonstrate that nuclear deformation is force dependent, however anisotropy is not. We also confirmed that SR displayed no clear dependence on the orientation of the nucleus with respect to the AFM cantilever (see Supplementary Fig. S1d,e). Importantly, tip geometry did not influence the observed anisotropy, as repeating the 10 nN experiment on 3T3s with a spherical (10 μm diameter) AFM tip also resulted in anisotropic nuclear expansion (SR = 0.959 ± 0.008, for n=26 cells). Moreover, anisotropic nuclear behaviour persists, as shown by performing the experiment over long durations (15 min). Strain measurements revealed that the nucleus reached 90-95% of its ultimate strain within 1 min of loading and resulted in significant strain anisotropy (Supplementary Fig. S2a-c).

### The effect of cytoskeletal inhibition on nuclear anisotropy

To determine the effect of the cytoskeleton on nuclear deformation, we made use of two very well-known anti-cytoskeletal drugs, Cytochalasin D (CytD) and Nocodazole (Noco), to selectively inhibit polymerization of actin filaments and microtubules (MTs), respectively^6,15,26,28,29^. Immunofluorescent images of fixed cells stained for actin, tubulin, and DNA demonstrate clear differences between untreated and treated cells (Fig. 2a-d). A combination of Noco and CytD (NocoCytD) was also used in some cases, which led to depolymerisation of both actin and MTs (Fig. 2d). Treated cells responded to 10 nN of force similarly to untreated cells, with the majority of the deformation occurring within seconds following loading (Fig. 2e-g). Area expansion was greatest for cells treated with NocoCytD (29.3 ± 3.8%) and CytD (25.3 ± 2.9%), both of which deformed significantly more than untreated cells (6.7 ± 0.7%). Cells treated with Noco only resulted in a marginal increase in nuclear expansion (8.6 ± 0.8%) following a 10 nN load. In the absence of loading, initial size (area) of 3T3 nuclei remained unchanged following treatment with the cytoskeletal inhibitors (Supplementary Table S1, P > 0.2, all treatments).

**Figure 2.**
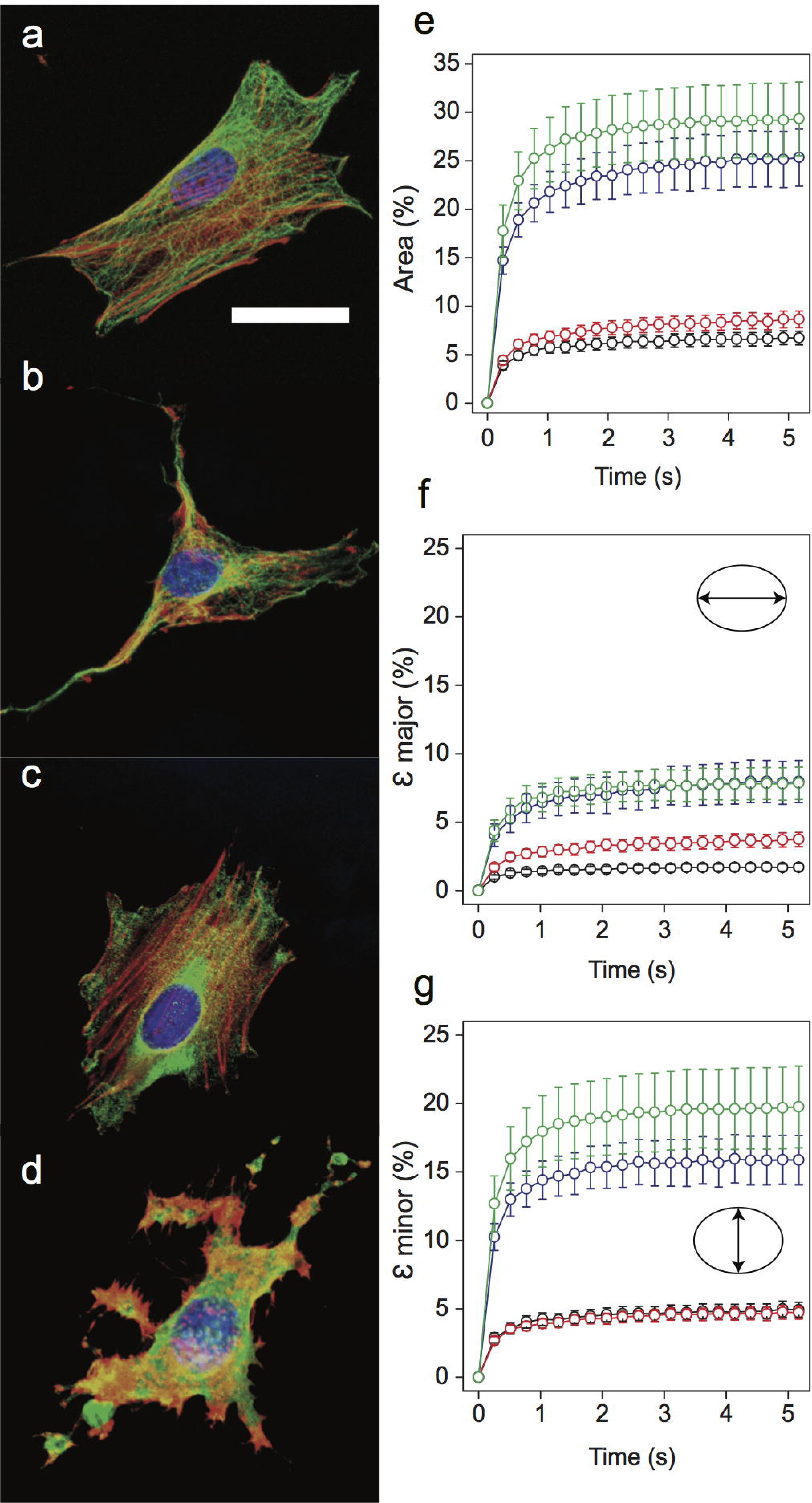
Cytoskeletal inhibition affects nuclear deformation. (a-d) Immunofluorescent images of fixed 3T3 cells showing DNA (blue), actin (red), and MTs (green). (a) Untreated 3T3 cells (n=32), (b) CytD (n=24), (c) Noco (n=24), and (d) NocoCytD (n=21) treated cells. Scale bar is 20 μm. (e) Change in projected nuclear area as a function of time for untreated (black), Noco (red), CytD (blue), and NocoCytD (green) treated 3T3s. Corresponding strains shown for (f) the major axis ( major) and (g) the minor axis ( minor) over 5 s of a 10 nN load. Shown is mean ± s.e.m.

In general, there is a statistically significant drug-dependent response in 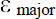 and 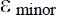 for cells treated with CytD and NocoCytD (Fig. 2f-g and Supplementary Table S1). Cells treated with NocoCytD exhibited the largest values of 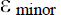, followed by cells treated exclusively with CytD. These two treatments also resulted in large strains in the major axis (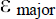). Surprisingly, in comparison to untreated cells, Noco resulted in a significant (P < 0.05) increase in 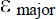. Following fits of the time dependent deformation to a Kelvin Voigt model, calculations of SR (Fig. 3a and Supplementary Table S1) revealed anisotropy in nuclear strain (SR < 1) that remains apparent irrespective of an intact cytoskeleton. Cells treated with CytD (alone or in combination with Noco) led to SR = 0.934 ± 0.016 and SR = 0.911 ± 0.020, respectively. Both treatments were statistically significant (P < 0.05) from untreated cells (SR = 0.971 ± 0.005), resulting in increased nuclear anisotropy. This loss of an intact actin network also significantly (P < 0.05) increased the deformation time constant in the minor nuclear axis (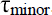) (see Supplementary Fig. S2d). In contrast, the loss of MTs alone (Noco) resulted in a significant increase in SR (0.990 ± 0.005, P < 0.01), indicating a near-isotropic nuclear deformation response. Noco treatment has been previously shown to activate RhoA^30^, resulting in increased actin stress fibre and focal adhesion formation. Here, stress fibres were apparent in the basal plane (Fig. 2c), but did not result in nuclear shape change, nor a decrease in minor axis strain (along the direction of fibres) (see Fig. 3b,c).

**Figure 3.**
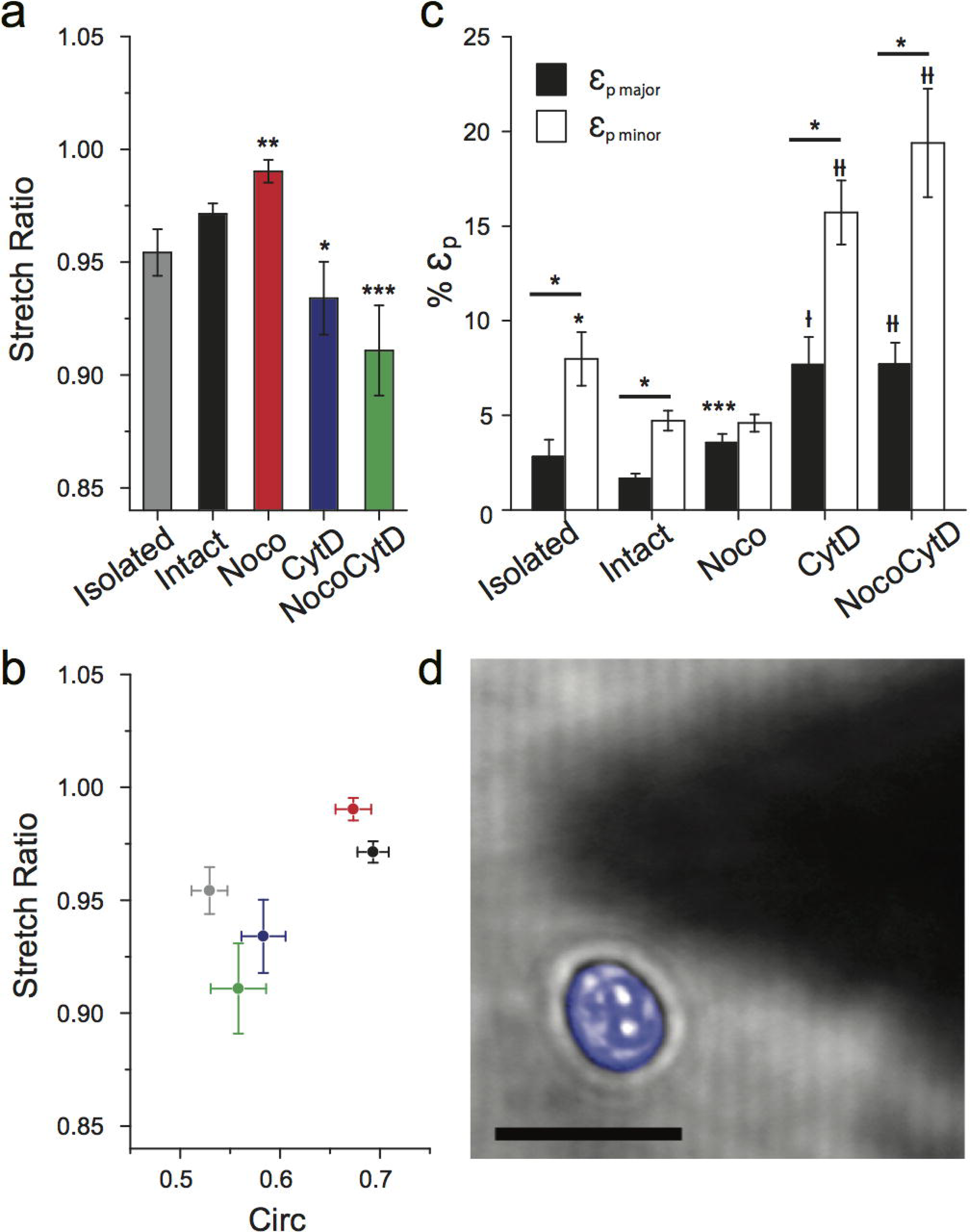
Nuclear anisotropy is intrinsic. (a) Mean anisotropic stretch ratio (SR) is shown for untreated and treated 3T3s. Anisotropy was demonstrated in all cases (SR < 1). Noco treatment demonstrated significantly less anisotropic behavior, whereas 3T3’s treated with either CytD or NocoCytD resulted in significantly more anisotropic behavior than untreated (intact) cells (* P< 0.05, ** P < 0.01, *** P< 0.001, with t-test). (b) Plot of anisotropic stretch ratio versus circularity. Nuclear shape (prior to deformation) is significantly less circular (more elliptical) for cells lacking an intact actin network. Untreated 3T3 (black), Noco (red), CytD (blue), NocoCytD (green), and isolated nuclei (grey). Shown is mean ± s.e.m. (c) Mean plateau strains (ε_p_) from fits to Kelvin-Voigt model. All cell populations, except those treated with Noco, exhibited significantly greater strain in their minor axes relative to their major nuclear axes (P < 0.05, with paired t-test). Shown is significance with respect to untreated (intact) cells (* P< 0.05, *** P< 0.001, † P < 0.0001, †† P < 0.00001, with t-test). (d) Typical isolated 3T3 nucleus (blue). Forces applied to elongated isolated nuclei (n=23) by an AFM tip (shadow) demonstrated reduced SR (increased anisotropic nuclear deformation). Scale bar is 20 μm.

Nuclear circularity (Circ) was measured prior-to and in response to extracellular force (see Supplementary Methods). Results indicate that anisotropic strain induces an increase in circularity (the nucleus becomes less elliptical) due to the fact that 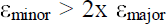 (Fig. 1d,e). Moreover, circularity drastically decreased with CytD and NocoCytD treatments, but was less pronounced for cells treated solely with Noco. In other words, upon loss of actin (alone or in combination with MTs), the nucleus became more elliptical. Force application also increased the circularity of all cells on average, as expected (Supplementary Fig. S3a). Although nuclear shape depends largely on the actin cytoskeleton, there is no clear correlation between nuclear shape and the observed level of anisotropy following loading (see Supplementary Fig. S3b). Cell cycle, as shown by a thymidine block of 3T3s, may be responsible for variability in nuclear shape, size, and deformability in untreated cells (Supplementary Fig. S3c,d).

Inside intact cells, actin clearly plays a major role in governing nuclear shape (in concert with the nucleoskeleton)^5,15,22^. Therefore, we isolated nuclei in order to probe its mechanical anisotropy in the absence of any confining structures (Fig. 3d). Results indicated that isolated nuclei became less circular (P < 1×10^−8^) after removal from the cell, suggestive of an intrinsic nuclear prestress. Isolated nuclei (103 ± 4 μm^2^) were smaller in cross-sectional area than intact cells (140 ± 5 μm^2^), correlating with cells having a depolymerized actin network. Following loading, average strain increased by nearly 1.7-fold in isolated nuclei in the major 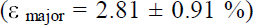 and minor axes 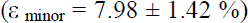, in comparison to nuclei deformed within intact cells. Importantly, isolated nuclei were also observed to deform anisotropically (SR = 0.954 ± 0.010, n=23), similar to nuclei within intact cells (SR = 0.971 ± 0.005, n=32). This result suggests that anisotropic mechanical properties are inherent to these nuclei, and likely depend on nuclear structure, possibly dictated by chromatin organization and/or key structural nuclear proteins (lamins).

### The role of nuclear architecture

Chromatin organization plays an important role in regulating transcription processes^31^ and governing the physical properties of the nucleus^15,32^. Moreover, extra-cellular forces impact the structure and organization of the nucleus and chromatin remodelling^7,8^. Here, to address the role of chromatin organization in anisotropy, we treated 3T3s with Trichostatin A (TSA), a specific inhibitor of histone deacetylase (HDAC), which leads to chromatin decondensation. TSA has been used in several studies of nuclear mechanics^33-35^. Cells treated with TSA underwent large morphological changes; they appeared more well-spread and exhibited extremely long extensions in comparison to untreated cells (Fig. 4a) consistent with previous reports^34,36^. In tune with the whole cell, nuclei of TSA-treated 3T3s (196 ± 13 μm^2^) were significantly (P < 0.05) larger in cross-sectional area than untreated nuclei (127 ± 5 μm^2^), (values reported here are from the same population). TSA treatment led to a significant decrease in strain in the minor nuclear axis, in response to an applied force (Fig. 4b), in comparison with untreated cells from the same population. This decrease in minor strain led to an overall decrease in the anisotropic response of TSA-treated 3T3 nuclei, as indicated by the anisotropic stretch ratio (SR = 0.980 ± 0.005), which was significantly (P < 0.05) greater than untreated cells (SR = 0.953 ± 0.006) (Fig. 4c). Nuclear circularity also decreased upon TSA treatment likely due to the elongation of the cells (P < 0.01) (Fig. 4c). These observations suggest two very important phenomena. First, disrupting chromatin structure directly impacts mechanosensitivity, as evidenced by the fact that cells displayed a reduction in the response to force in the minor axis. Second, TSA-treated nuclei can be characterized by a more isotropic deformation. While TSA has been shown to influence microtubule dynamics and focal adhesion turnover^37^, treatment also leads to clear histone deacetylation, as seen by pronounced nuclear expansion (correlating with cells blocked in G1/S phase, Fig. S3c,d). Chromatin organization clearly plays a critical role in regulating the mechanosensitive deformation response of the nucleus to mechanical forces arising in the extracellular environment, but may be inextricably linked to the effect on the cytoskeleton here.

**Figure 4.**
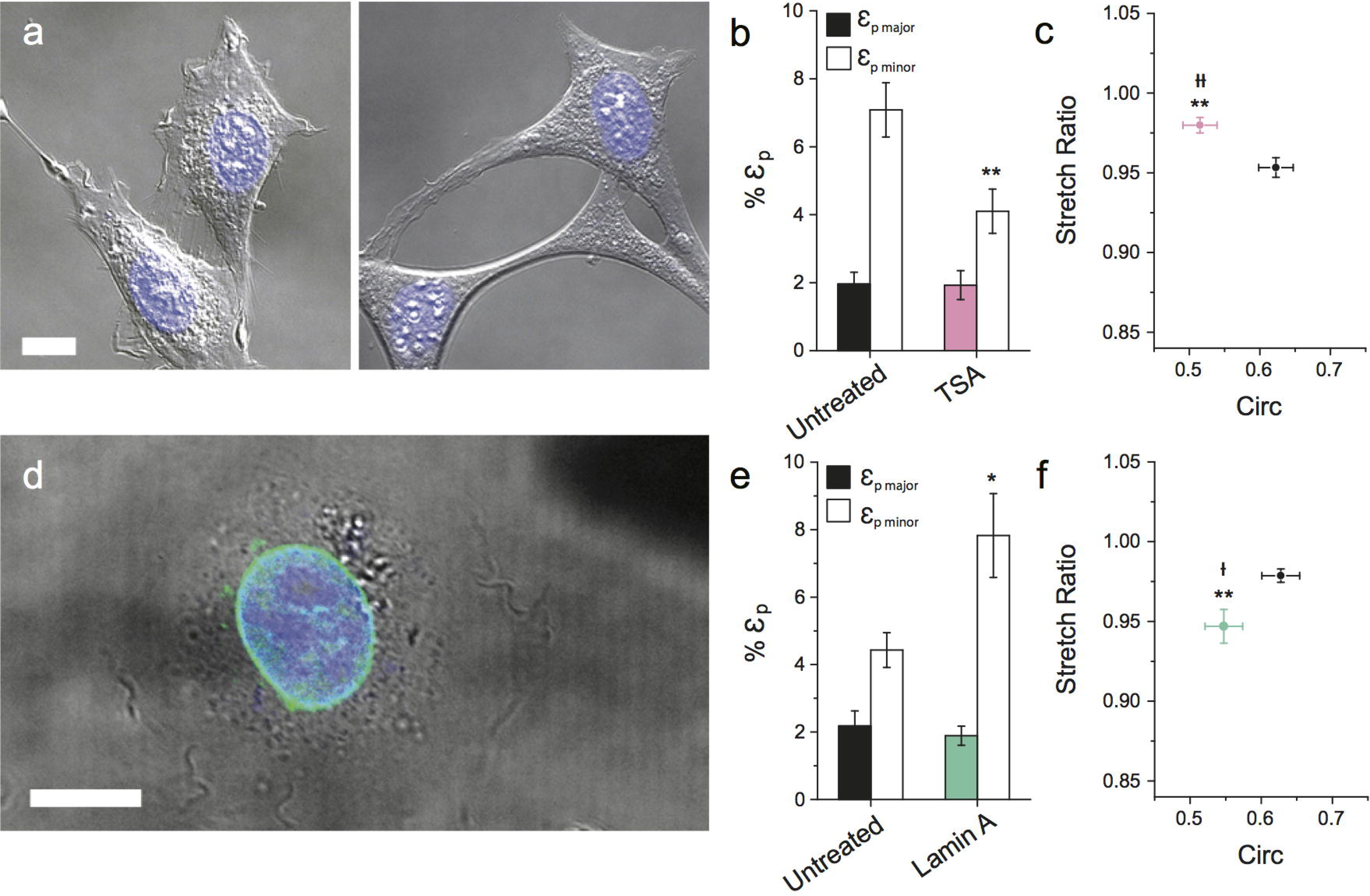
Role of nuclear architecture in anisotropy. (a) DIC images of live untreated 3T3 cells (n=30) (left) and cells treated with TSA (n=34) (right). Scale bar is 10 μm. (b) Plots of mean plateau strains (εp) are shown. (c) Plot of mean stretch ratio and circularity for untreated (black) and TSA-treated (pink) 3T3s. (d) Over¬expression of lamin-A in 3T3s, shown by live-cell image transiently expressing EGFP-lamin-A in the nuclear envelope. DNA in blue, lamin-A EGFP (green). Scale bar is 12.5 μm. (e) Plots of mean plateau strains (εp) are shown for untreated (black) and Lamin-A over-expressing 3T3s (green). (f) Plot of stretch ratio and circularity for untreated (n=22) and lamin-A transfected cells (green) (n=18). (* P< 0.05, ** P< 0.01, with t-test for strains and stretch ratios, † P < 0.05, †† P < 0.01, with t-test for circularity).

Alternatively, lamin proteins are known to influence nuclear mechanics and structure. Lamin deficient mice have been employed in previous studies of nuclear mechanics^9,22,34,38^. Complications can arise in this approach due to numerous altered states and compensation mechanisms that occur in response to lamin deficiency^9,38^. Therefore, we employed the alternative strategy of transiently over-expressing EGFP-lamin-A in 3T3 cells. EGFP-lamin-A was shown to localize at the nuclear membrane (Fig. 4d). Surprisingly, lamin-A over-expression resulted in a significant increase in strain in the minor axis 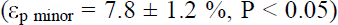, in comparison to untreated 3T3s from the same population 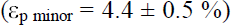. This increase in strain along the minor axis resulted in increased anisotropic nuclear deformations (P < 0.01) (SR = 0.947 ± 0.011) for cells over-expressing lamin-A, in comparison to wild type (SR = 0.979 ± 0.004). Lamin-A over-expression also resulted in a significant decrease in circularity of 3T3s (P < 0.05). These results suggest that lamin-A, in concert with chromatin and cytoskeletal organization, influences the level of mechanical anisotropy observed in nuclei in response to extracellular forces.

### Widespread observations of nuclear mechanical anisotropy

To examine whether nuclear anisotropy is a pervasive phenomenon, we performed the short deformation experiment (10 nN for 5 s) on a variety of other cells (immortalized and primary fibroblasts, epithelial, myoblast and embryonic stem cells) derived from several species (human, mouse, canine, hamster). Although strain magnitudes varied widely in response to force (Supplementary Table S2), in all cases the mean SR was observed to be < 1 (Fig. 5). Nuclear shape, as characterized by circularity, varied greatly between cells and again did not correlate with SR. Importantly, strain in the minor axis was significantly (P < 0.05) greater than strain in the major axis for all cells examined, except for mESC and cells cultured from Dmd^mdx^ mice. Overall, these results demonstrate that deformation anisotropy appears to persist among the diverse cell and species types surveyed here.

**Figure 5.**
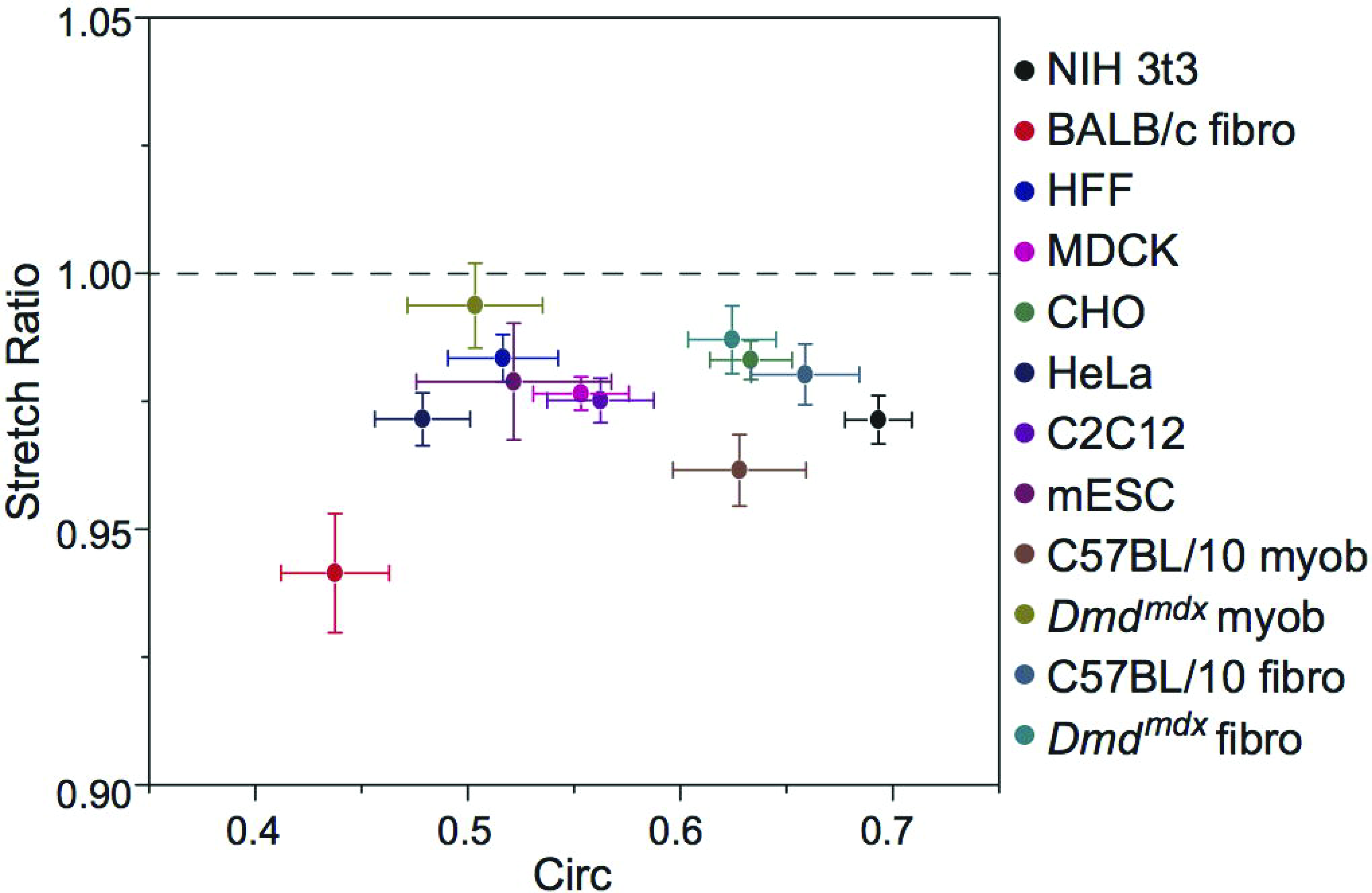
Nuclear anisotropy is consistent across cell types. Plot of anisotropic stretch ratio (SR) versus circularity for various established cell lines: 3T3s (n=32), HFF (n=32), CHO (n=17), MDCK (n=30), HeLa (n=26), C2C12 (n=20), D3 mESC (n=10), and primary cells including fibroblasts from BALB/c mice (n=15), and fibroblasts from C57BL/10 (n=30) and Dmd^mdx^ (n=22) mice, as well as myoblasts from C57BL/10 (n=16) and Dmd^mdx^ (n=9) mice. Established cell types demonstrated a clear nuclear anisotropy, as demonstrated by SR consistently < 1. Deformations associated with Dmd cell types were more isotropic than wild-type cells, possibly due to a lack of lamins A/C and altered chromatin organization. Shown is mean ± s.e.m.

## Discussion

Observations of nuclear anisotropy^15,22,23^, and recent auxetic behaviour in ESC^39^, suggest that the mechanical properties of the nucleus are not isotropic, as often described^9,14,20,24,25^. Here, we examined whether anisotropic nuclear mechanics could be observed within living cells when exposed to locally applied nano-forces. This is of crucial importance, as localized extracellular forces may impact gene regulation and signalling through direct physical changes in nuclear and genomic structure^7,8^. Therefore, systematically characterizing the deformation of the nucleus in response to force, and its mechanical anisotropy, is a critical first step in understanding this complex process. Here, using 3T3 fibroblasts as a model, we demonstrated that nuclei deform anisotropically in response to externally applied point-forces from an AFM tip. The deformation occurred very rapidly (< 1 s); eventually reaching a plateau that persists for long periods (up to 15 min, max length of study). Strain along the minor nuclear axis was ∼50% larger than the strain along the major axis. This anisotropy was characterized by a stretch ratio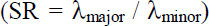, with mean values of SR < 1, consistent with recent studies^15,22,23^. Importantly, our results demonstrate that controlled anisotropic deformation of the nucleus occurs in direct response to both point-forces (from a pyramidal tip) and large-contact extracellular forces (from a spherical tip) (Supplementary Table S1).

Using 3T3s, we set out to characterize the origins of nuclear anisotropy. With the use of specific inhibitors, cytoskeletal elements were shown to play important roles in regulating nuclear shape, as well as the magnitude, timescale, and anisotropy of nuclear deformation (Fig. 3). The actin network has a profound impact on nuclear shape, demonstrated by significant reductions in circularity upon its depolymerization. A loss of intact actin also caused a significant increase in the magnitude of nuclear deformation along both axes, as well as the rate of expansion along the minor axis (Fig. S2). Actin is highly cross-linked to the nuclear architecture through nesprin-1 and -2 proteins^3,5^ and clearly plays an important role in regulating nuclear deformations. Actin also contributes significantly to the regulation of nuclear anisotropy, consistent with previous studies^15,23^. Importantly, a loss of an intact network of MTs resulted in the opposite effect. Increased strain was observed along the major axis, which led to an increase in SR (Fig. 3). MTs have been shown to distribute radially near centrosomes^40^, and are now well known to contribute to cell polarization during migration independently of actin^41^. It is plausible that perinuclear MTs create a dense filament mesh at the ends of the major axis, contributing to strain resistance and the observed anisotropy. These results correspond with our other recent findings suggesting an MT-dependent resistance in the major axis, and actin-dependent resistance in the minor nuclear axis, following strain via planar stretch^23^.

Finally, isolated nuclei revealed that anisotropic deformation is intrinsic to the structure of the nucleus itself. Contrary to previous reports^22^, in the complete absence of prestress or confinement arising from the cytoskeleton/cytoplasm, isolated nuclei display strain that primarily occurs along the minor axis resulting in anisotropic deformation. It appears that the cytoskeleton is designed to regulate the degree of nuclear anisotropy in 3T3s (Fig. 6). MTs resist nuclear deformations in the major axis, while actin resists deformations in the minor axis. The respective cytoskeletal networks act as effective springs that resist deformation of the nucleus against external (apically applied here) forces. This interplay between the cytoskeletal networks limits nuclear deformations, yet permits the inherent anisotropic mechanics observed in isolated nuclei. Moreover, our results suggest that an outward force acts on intact nuclei by the cytoskeleton (due to adhesion), as isolated nuclei were much smaller in size. It has been proposed that anisotropy in global cellular deformation is dominated by the presence of actin fibres, which tend to align parallel to the long axis of the cell^42^. Our results strongly agree with these findings, as the major axis tended to align with the polarization of the actin cytoskeleton (seen in our previous work^6^). While actin plays a key role in dictating nuclear shape and its resistance to deformation, nuclear anisotropy appears to initially stem from the existence of an intrinsically prestressed nucleus due to the nucleoskeleton, nuclear envelope and/or chromatin organization^3,5^.

**Figure 6.**
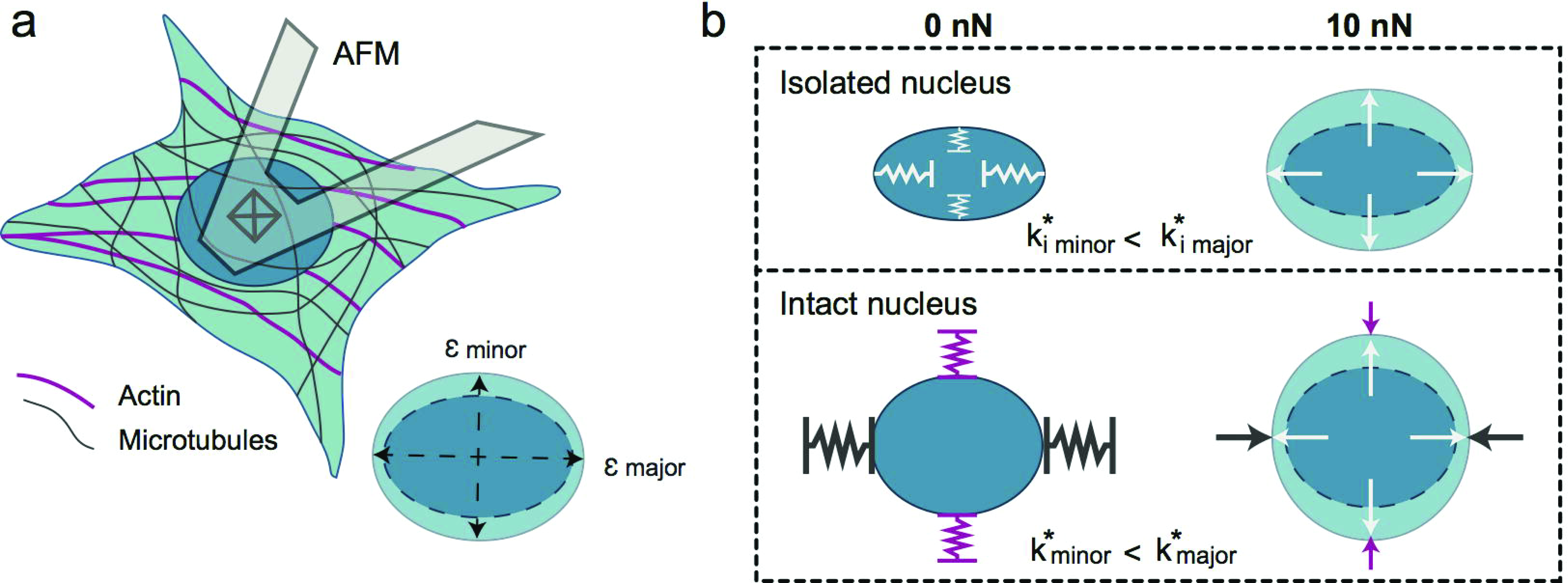
Intrinsic nuclear anisotropy is regulated by cytoskeletal resistance. (a) Shown is a schematic of a 3T3 cell deformed by an AFM tip, with an enlarged depiction of the resultant nuclear strain along the major and minor axes. (b) Depicted is the nuclear response to force in relation to the observed mechanics. Anisotropic strain occurred in isolated nuclei, corresponding to a larger resistance to deformation in the major axis, opposed to its minor axis (represented by effective internal spring constants, k_i_^*^, in the minor and major axes). For an intact cell, nuclear deformations are regulated by the cytoskeleton. In response to an apically applied force, actin resists deformations in the minor axis, while microtubules resist strain along the major axis. Resistance to deformation along the major axis is greater than the minor axis, as shown by effective spring constants k^*^, allowing the cytoskeleton to promote the inherent anisotropy.

Both chromatin organization and lamins have been shown to dictate nuclear mechanics through their coupled behavior^43^, and so were hypothesized to regulate nuclear anisotropy. First, TSA was used as an HDAC inhibitor to de-condense chromatin in 3T3s^33-35^. TSA treatment resulted in large morphological changes, likely due to the inhibition of gelatinase A, a matrix metalloproteinase^36^. Importantly, chromatin de-condensation resulted in a significant increase in nuclear size and demonstrated a significant impact on the mechanosensitivity of 3T3 nuclei (in agreement with previous work^44^); in particular, reduced minor axis strain. Importantly, TSA treatment diminished the anisotropic response (Fig. 4). However, future studies are required to decouple the cytoskeletal side effects of TSA treatment from chromatin decondensation. In contrast, over-expression of lamin-A led to an increase in strain along the minor nuclear axis, resulting in an increased anisotropic response (Fig. 4c). The increase in minor axis strain was surprising considering that lamin-A is known to provide structural stability to the nucleus^9,16^. However, nuclear anisotropy associated with lamin-A expression is in line with previous observations wherein lamin-A/C deficient MEFs demonstrated isotropic nuclear deformations, in contrast to wild type MEFs^22^. Localization of lamin-A in polarized 3T3 nuclei may provide stability along the ends of the nucleus, where it has been suggested that the majority of nucleo-cytoplasmic contacts can be found^15^. The importance of lamins should not be underestimated, since they influence overall nuclear stiffness and play an important role in chromatin regulation^45^. Nuclear mechanical anisotropy may indeed arise from a force balance between chromatin organization (outward entropic force) and lamin-A localization (a containment force), which could account for the inherent non-isotropic mechanical properties (Fig. 6). Nuclear organization is certainly tied to cell cycle^46^, as are its mechanics. For example, the expression of lamin-A has been linked to differentiation^47^, and recently lamina-associated chromatin organization has been shown to remodel during mitosis^48^. Here, blocking 3T3s in the G1/S and G2/M transition phases resulted in a significant reduction in nuclear circularity (Fig. S3d) but increased nuclear area (Supplementary Table S3, P < 0.05). Importantly, cell cycle was shown to have a significant effect on nuclear deformability (Fig. S3), and can account for variations observed across populations of untreated cells. Implications of cell cycle in nuclear anisotropy warrants further investigation, particularly into the coupled effects of nuclear organization and structural mechanics throughout the progression of interphase. Although speculative, regulated polarization of nuclear mechanics in this way may play a role in force-mediated transcription processes^31^.

Nuclear deformation anisotropy is widespread, as demonstrated by performing the experiment on a variety of established cell and species types. Although nuclear shape and strain magnitudes following perturbation varied widely, SR was always observed to be < 1 (Fig. 5). This survey showed that highly circular nuclei displayed anisotropic deformations, and that initial nuclear shape did not influence anisotropy. Interestingly, diseased primary cells (Dmd^mdx^ myoblasts and fibroblasts) resulted in increased strain in the major axis in comparison to cells isolated from control mice (C57BL/10) (Supplementary Table S2). This led to a significant increase (P < 0.05) in SR for Dmd^mdx^ myoblasts in comparison to their wild-type control, indicating a near-isotropic response. A reduction in anisotropy and change in major axis strain may be attributed to global histone modification, as witnessed by dispersed and open chromatin configurations in Dmd^mdx^ myoblasts^49,50^. Importantly, pluripotent mESCs also displayed anisotropic nuclear deformations. However, 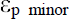 was only marginally greater than 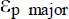 (P > 0.05, paired t-test), a result likely influenced by lamin-A deficiency^22^, as observed in these cells (Fig. S4). Recently, mESCs in a transitional state were shown to exhibit complex auxetic behaviours in response to compression^39^. Moreover, mESC in a naïve pluripotency state responded with auxetic behaviour only upon TSA treatment^39^. Those findings suggest that mESC mechanical properties are highly complex and depend on transitional state and chromatin organization. Taken together, these results correspond with nuclear mechanics being largely dependent on the coupling of chromatin organization and lamin-A expression. Nonetheless, anisotropy appears to be a highly conserved nuclear characteristic, at least among the diverse cell types examined here.

Although we have not established the functional role of nuclear anisotropy, it is intriguing that 3T3 fibroblasts exhibit such a high degree of regulation over nuclear deformation. Moreover, the observed anisotropy was conserved among diverse cell types from several species, suggesting that the nucleus should be treated as an anisotropic, not isotropic, material. Therefore, future studies will focus on examining how key proteins (post-translational modifiers to histone tails^51^) involved in chromatin organization, chromatin territories and the nucleoskeleton regulate deformation anisotropy^5,31,52^. Moreover, several studies have shown that cyclic stretch can induce gene expression changes and it remains unclear how inhibiting anisotropy (TSA treatment or specific knock outs) may impact these findings^32,53,54^. Clearly, more work is required to confirm whether or not nuclear deformation anisotropy is simply a result of nuclear organization, or if it plays a functional role in cell biology. Regardless, we have shown that living 3T3 cells regulate nuclear deformation at the cytoskeletal and organizational levels of the nucleus itself, in such a way that the nucleus is significantly more sensitive to deformation along the minor axis. Notably, anisotropic deformation occurs whether the mechanical stress is emanating from below the cell (substrate stretch^23^) or from above (direct AFM compression), or whether locally or globally applied. Though speculative, we suggest that nuclear structure, organization and material properties are optimized to enhance its sensitivity to extra-cellular forces in order to promote mechanically induced gene expression.

## Methods

Detailed descriptions of reagents, cell lines, animal models, plasmids, simultaneous AFM-LSCM, image analysis and statistical tests are contained in the Supplementary Methods. All experimental procedures involving laboratory animals were approved by the Animal Care and Use Committee of the University of Ottawa. All experiments were carried out in accordance with the approved guidelines. All data presented are mean ± s.e.m. Statistical significance is considered for P-values where P < 0.05, as determined by two-tailed student’s t-tests, unless noted otherwise.

## Acknowledgments

The authors acknowledge support from individual Natural Sciences and Engineering Resource Council (NSERC) Discovery Grants (J.L.H. and A.E.P.) and an NSERC Discovery Accelerator Supplement (A.E.P.). C.M.C. was supported by a postdoctoral training award from the “Fonds de Recherche du Québec - Santé” (FRQS). A.E.P also acknowledges generous support from the Canada Research Chairs program and a Province of Ontario Early Researcher Award. The authors would like to thank Daniel Modulevsky for advice on experimental procedures and Adriana Prystay for culture of CHO cells.

## Author contributions

K.H., J.K.M. and A.E.P. designed the research; K.H. performed the research with some contributions from J.K.M., C.E. and A.E.P.; K.H. and A.E.P. analyzed the data; C.M.C., S.H. and I.S.S. supplied primary and mESC cells, respectively; J.L.H. contributed analytical methods; K.H., J.K.M. and A.E.P. wrote the paper.

